# Prospective evaluation of renal function in dogs with chronic mitral valve disease

**DOI:** 10.1101/2020.03.18.996785

**Authors:** Elisa Martinelli, Serena Crosara, Chiara Locatelli, Anna Maria Zanaboni, Paola Brambilla, Cecilia Quintavalla

## Abstract

The coexistence of renal and cardiac disease has been defined in dogs and cats as cardiovascular-renal disorders (CvRD). In humans, renal function is affected by recurrent episodes of acute congestive heart failure (CHF). The aim of this prospective, case-control study was to evaluate the appearance and influence of worsening cardiac disease (WCD), defined on echocardiographic and radiographic parameters, on renal function (defined as worsening renal function [WRF], on the basis of serum creatinine level and presence of proteinuria) in two population: 21 dogs with chronic mitral valve disease (CMVD) and 20 healthy dogs. Dogs were sorted into groups according to the presence/absence of WRF or WCD. Statistical analysis was performed between CMVD dogs and healthy dogs and inside the CMVD dogs group. There was no statistically significant difference in developing WRF between dogs with/without WCD and no statistical evidence to support a difference in WRF parameters in dogs experiencing CHF and dogs not experiencing it. The prevalence of azotemia in CMVD dogs was significantly higher than the prevalence of azotemia previously reported in the general population of dogs. Diuretics therapy didn’t affect renal function. No difference in survival time was seen between groups. In conclusion, CHF, WCD and diuretics therapy didn’t directly induce WRF. However, considering the prevalence of azotemia, data suggests a link between heart and kidney function (despite we didn’t excluded aged-related coexistence of organ damage). A bigger number of dogs at inclusion is required to reach statistical significance.

## Introduction

The heart and kidney are both involved in basic physiology, and their functions are strictly linked, that’s why primary disorders of heart or kidney often result in secondary dysfunction or injury to the other organ [1–5]. The coexistence of renal and cardiac disease, referred as cardiorenal syndrome (CRS) in human medicine and as cardiovascular-renal disorders (CvRD) in veterinary medicine, significantly increases mortality and morbidity in human patients [6]. Cardiorenal syndrome was first described in 1951, however, the existence and importance of CvRD in dogs is still unknown [7–12]. The most common heart disease affecting dogs and leading to congestive heart failure (CHF) is chronic mitral valve disease (CMVD) while, in humans, coronary artery disease and systemic hypertension are most frequent [13–15]. Recurrent episodes of acute heart failure are considered one of the causes leading to worsening renal functions (WRF) in human medicine [16]. The aim of our study was to assess the influence of CHF and/or worsening of cardiac disease (WCD), defined on echocardiographic and radiographic parameters, on renal function, through the evaluation of elected parameters (Table 1).

**Table 1.**
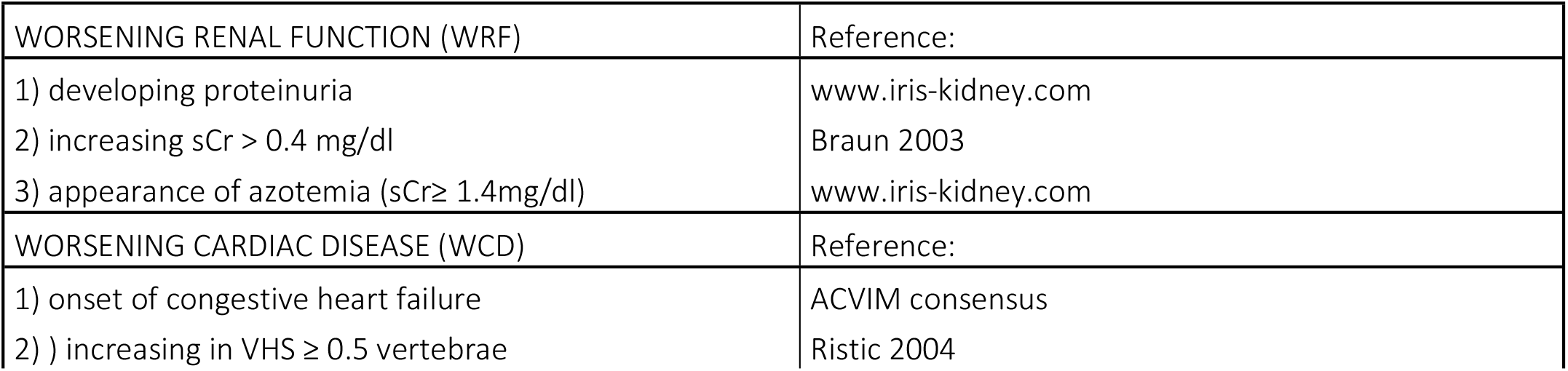
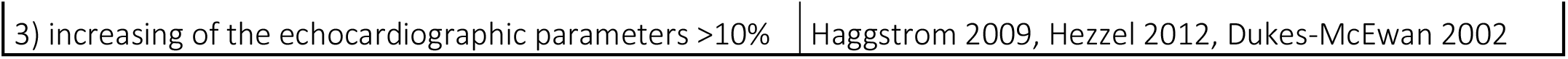
Parameters considered for worsening renal function (WRF) and worsening cardiac disease (WCD).

## Materials and Methods

This double-center, prospective, case-control study was conducted at two Veterinary Teaching Hospitals in Italy, in accordance with the principles of Good Clinical Practice (Directive 81/852/EEC as amended; DIRECTIVE 2004/28/EC as amended). Consent for each dog was obtained from the owners before the enrolment. Permission to conduct the study was received from the Ethics Committee of both the University involved (Protocol number 22/2013).

Dogs included were recruited among those examined during routine clinical practice, between January 2013 and May 2013. Information obtained from the medical records included signalment and history. All the included dogs (cases and controls) underwent physical examination, thoracic radiographs, electrocardiography (ECG), complete blood count (CBC), serum biochemical analysis, complete urine analysis, urine protein/creatinine ratio (UPC) and indirect systemic blood pressure evaluation by Doppler flow meter [17]. Dogs were re-evaluated every 6 months until October 2014 and data were collected in a dedicated datasheet. At the end of the study, dogs were sorted into groups according to the presence/absence of signs of WRF and WCD defined at 12 month after the inclusion date to perform statistical analysis.

The inclusion criteria in the CMVD group were: dogs firstly diagnosed with CMVD (CMVD dogs) classified as ACVIM class B2 (asymptomatic patients with hemodynamically significant valve regurgitation associated to mitral leaflet thickening, prolapse or both, and echocardiographic evidence of left-sided heart enlargement), or dogs in class ACVIM C at their first occurrence of clinical signs of CHF. The left heart was considered remodeled when the left atrial–to-aortic root ratio (LA/Ao) was ≥ 1.6 and the normalized left ventricular end-diastolic diameter calculated according to Cornell’s method of allometric scaling (LVEDDn) was ≥ 1.7 [18, 19].

The exclusion criteria were: other congenital or acquired heart disease, azotemia (sCr ≥ 1.4 mg/dl), neoplasm, systemic or metabolic disease (hyperadrenocorticism, hypothyroidism, diabetes mellitus, lower urinary tract disease), hypertension or hypotension (defined as SBP≥160 mmHg and <80 mmHg respectively) [17]. Dogs in treatment for CHF, working dogs or dogs receiving high proteins diets were excluded [20].

Healthy adult dogs, older than 6 years, selected during screening examinations for the medical status of senior pets, performed at the authors’ institutions according to American Animal Hospital Association (AAHA) Senior Care Guidelines for Dogs and Cats, were included in the control group [21–23]. All dogs were determined to be healthy on the basis of thorough physical and cardiovascular examinations, including ECG, echocardiography, thoracic radiographs and systolic arterial blood pressure measurement. Exclusion criteria were: dogs under medications known to affect the cardiovascular, renal or respiratory systems; dogs with uncooperative temperament that might require sedation for an echocardiogram; the evidence of arrhythmias on ECG except sinus arrhythmia; cardiac abnormalities and a SBP≥160 mmHg.

All thoracic radiographs were acquired at the peak of inspiration. For each set of thoracic radiographs, the vertebral heart score (VHS) and the pulmonary patterns (interstitial, alveolar or bronchial) were evaluated by two operators at each center (E.M. and C.Q. or E.M and C.L.) in order to estimate heart size and presence of CHF respectively [24]. Normal heart size was defined as VHS ≤ 10.5 on right lateral view of the thorax [19–25]. All the radiographic measurements were performed three times and averaged. Cardiogenic pulmonary edema was defined when pulmonary venous congestion and/or an interstitial or alveolar lung pattern were associated with cardiomegaly and clinical signs consistent with left-sided CHF including increased respiratory rate [26].

All echocardiographic studies were performed using an Esaote MyLab50 ultrasound machine (Esaote Medical System), equipped with multi-frequency phased array transducers (7.5-10 MHz and 2.5-3 MHz). All echocardiographic measurements were made on conscious dogs, in accordance with the guidelines of the American Society of Echocardiography, by trained observers (E.M., C.L., C.Q.). Interventricular septal thickness (IVS), left ventricular internal diameter (LVID), and left ventricular posterior wall thickness (LVPW) in diastole (d) and systole (s) were obtained from the right parasternal short-axis M-mode view at the chordae level using the leading-edge to leading-edge method [27]. Aortic root diameter (Ao) and left atrial diameter (LA) were obtained from 2D right parasternal short-axis view using the Hansson’s method [28]. Mitral valve inflow (E peak velocity - Evmax, A peak velocity – Avmax, E/A ratio), and peak velocity of mitral and tricuspid regurgitations (MR and TR) were evaluated using the pulsed and continuous spectral Doppler under color Doppler guidance [29]. Variables calculated were: LA/Ao, left ventricle end-systolic volume index (ESVI), left ventricle end-diastolic volume index (EDVI), LVEDDn, left ventricular fractional shortening (FS%) and ejection fraction (EF%) [18]. The EDVI and ESVI were calculated according to the Teichholz formula and normalized to body surface area (BSA).

A 5 minutes standard 6-lead ECG was obtained in awake dogs to assess heart rate and rule out cardiac arrhythmias that could affect haemodynamics and renal perfusion

A venous blood sample (approximately 3 ml) was collected from animals fasted at least for 12 h. Serum was separated and stored at 4°C and processed within 24 hours. Haematology was performed using a laser hematology analyser (Sysmex XT-2000iV, Sysmex; Cell-Dyn 3500 Plus, Abbott), equipped with a multispecies software. Quality control and calibration were periodically performed with e-check Xe (Sysmex). The following parameters were recorded: white blood cell count (WBC), red blood cell count (RBC), hemoglobin (Hb), hematocrit (Ht), mean corpuscular volume (MCV), mean corpuscular hemoglobin (MCH), mean corpuscular hemoglobin concentration (MCHC). Biochemistry was performed using an automated spectrophotometer (Cobas Mira, Roche; Cobas Integra Plus, Roche) with reagents provided by Real Time. The following analytes were measured: glucose (GLY; GOD-POD method), serum urea (UREA; Urease method), sCr (modified Jaffe’ method) and total proteins (TP).

Urine sample were collected by cystocentesis or free catch. Five milliliters of each sample were centrifuged for 5 min at 500 g; then the supernatant was frozen at −20 °C into a plain tube. The remaining urine was used to resuspend the urine sediment that was examined microscopically at 400× magnification to count the mean number of red blood cells and white blood cells per high power field. The presence of epithelial cells, casts, crystals, bacteriuria, spermaturia and lipiduria were evaluated according to a semiquantitative scale (rare, moderate, abundant, or very abundant). Calculation of the UPC was performed in batches, after a maximum storage of 2 days as follow: urinary proteins were measured using pyrogallol red (Total Proteins High Sensitivity, Ben Biochemical Enterprise) and urinary creatinine was measured with a modified Jaffe method (Real Time Diagnostic Systems). Samples were manually diluted 1:20 with distilled water to fit the linearity of the method. Occasionally, particularly concentrated urine samples were further diluted to 1:100 to fit the linearity of the method.

### Statistical methods

Statistical analysis was performed using IBM SPSS Statistics 20 [Release 2.07 GPL Edition]. Normality of the distribution was tested using non-parametric Shapiro-Wilk tests. Student t tests and Mann-Whitney U Test were used to investigate differences between sets of data. Variables normally distributed were presented as mean and standard deviation (SD), whereas variables non-normally distributed were presented as median and interquartile (IQ) range. Correlation between variables was tested by the Pearson correlation coefficient. A p value <0.05 was considered significant. The considered variables were: parameters of renal function (sCr and UPC); parameters related to cardiac disease (VHS, the echocardiographic parameters LA/Ao, LVEDDn, EDVI, ESVI; furosemide’s dose in mg/kg/die and number of CHF episodes). Worsening cardiac disease and renal function were considered if at least one of the sentence reported in Tab 1 were confirmed.

A single investigator (E.M.) conducted telephone interviews with dog owners to determine the clinical outcome of each dog. Question asked to determine the clinical outcome were if the dog was dead or alive, if the dog had been euthanized or died spontaneously and which was the reason for euthanasia or the cause of death. Cardiac-related death was defined as death occurring because of progression of clinical signs of CHF. Euthanasia because of refractory CHF was scored as cardiac-related death. Sudden death was regarded as cardiac-related if no other cause of death was identified. Renal-related death was defined as death occurring because of progression of clinical signs of chronic kidney disease (CKD). Euthanasia because of refractory renal failure was scored as renal-related death. Survival time was counted from the day of inclusion in our study to either the day of death or closing time of the study (October, 2014). The end-points of the study were: all-cause death, cardiac related death and renal related death. Dogs still alive at the end of the study period were right-censored.

## Results

Twenty-one dogs affected by CMVD and 20 healthy dogs were included. Population characteristics are reported in Table 2. Breeds included in the CMVD group were mongrel (n = 12), miniature Poodle (n = 3), Bolognese (n = 1), Cavalier King Charles Spaniel (n = 1), Dachshund (n = 1), Deutsch Kurzhaar (n = 1), Doberman pinscher (n = 1) and Yorkshire terrier (n = 1). Breeds included in the healthy group were mongrel (n = 13), Pitbull (n = 4), Deutsch Kurzhaar (n = 1), Galgo espanol (n = 2). Laboratory variables, echocardiographic values and statistically significant difference between the two groups at inclusion are reported in Table 2 and 3 respectively.

**Table 2.** Population characteristics and laboratory variables.

| Variable | Overall |  | CMVD dogs |  | Healthy dogs |  | p |
| --- | --- | --- | --- | --- | --- | --- | --- |
|  | mean or<br>median | st.dev or<br>IQ range | mean or<br>median | st.dev or<br>IQ range | mean or<br>median | st.dev or<br>IQ range |  |
| No. of dogs | 41 |  | 21 |  | 20 |  |  |
| % male | 58 |  | 57 |  | 60 |  |  |
| BW (kg)** | 13.00 | 16.65 | 8.0 | 3.15 | 24.5 | 18.5 | p<0.01 |
| Age (years)* | 10.34 | 3.00 | 11.2 | 2.6 | 9.35 | 3.1 | p<0.05 |
| WBC (x 10<sup>3</sup> /μL) | 8.89 | 3.34 | 8.24 | 2.24 | 9.58 | 4.16 | NS |
| RBC (x 10<sup>6</sup> /μL) | 6.97 | 0.95 | 7.10 | 0.67 | 6.75 | 1.17 | NS |
| Hb (g/dl) | 15.72 | 1.71 | 16.17 | 1.53 | 15.26 | 1.81 | NS |
| Ht (%)* | 44.09 | 5.72 | 45.99 | 3.98 | 42.09 | 6.63 | p<0.05 |
| MCV (μ<sup>3</sup>) | 66.05 | 3.78 | 65.40 | 3.34 | 66.74 | 4.17 | NS |
| MCH (pg) | 25.34 | 10.08 | 22.94 | 1.03 | 27.86 | 14.13 | NS |
| MCHC (g/dl) | 35.00 | 2.45 | 34.90 | 5.10 | 35.10 | 2.17 | NS |
| UREA (mg/dl) | 32.20 | 13.55 | 33.00 | 7.00 | 31.9 | 9.90 | NS |
| sCr (mg/dl) | 0.75 | 0.23 | 0.78 | 0.23 | 0.73 | 0.23 | NS |
| GLY (mg/dl) | 85 | 13 | 93 | 8 | 77 | 14 | NS |
| TP (g/dl) | 6.47 | 0.58 | 6.48 | 0.62 | 6.46 | 0.56 | NS |
| UPC | 0.26 | 0.26 | 0.27 | 0.27 | 0.24 | 0.25 | NS |
| USG | 1029 | 11 | 1031 | 12 | 1027 | 10 | NS |

**Tab 3.** Echocardiographic values and statistically significant difference between the two groups at inclusion.

|  | Overall |  | CMVD dogs |  | Healthy dogs |  |  |
| --- | --- | --- | --- | --- | --- | --- | --- |
|  | mean or<br>median | st.dev or<br>IQ range | mean or<br>median | st.dev or<br>IQ range | mean or<br>median | st.dev or<br>IQ range | p |
| HR (bpm) | 116 | 40 | 120 | 23 | 121 | 29.9 | NS |
| LVIDd (mm) | 37.13 | 8.33 | 36.25 | 7.35 | 38.09 | 9.40 | NS |
| LVIDs (mm) | 20.25 | 8.18 | 18.40 | 2.70 | 24.30 | 10.70 | NS |
| LVEDDn | 1.72 | 0.34 | 1.94 | 0.20 | 1.50 | 0.30 | NS |
| EDVI (ml/m<sup>2</sup>) | 108.14 | 42.99 | 135.61 | 32.03 | 79.30 | 33.17 | NS |
| ESVI (ml/m<sup>2</sup>) | 27.72 | 14.64 | 30.28 | 13.72 | 25.03 | 15.43 | NS |
| EF%** | 80.16 | 9.67 | 84.18 | 4.99 | 75.72 | 11.63 | p<0.01 |
| FS%** | 43.00 | 8.76 | 46.49 | 5.69 | 39.14 | 10.02 | p<0.01 |
| LA (mm) | 25.96 | 5.92 | 25.09 | 6.01 | 27.10 | 5.81 | NS |
| Ao (mm)** | 17.79 | 4.91 | 15.00 | 3.36 | 21.46 | 4.17 | p<0.01 |
| LA/Ao** | 1.50 | 0.27 | 1.68 | 0.17 | 1.27 | 0.17 | p<0.01 |
| Evmax (m/s)** | 0.87 | 0.24 | 0.98 | 0.23 | 0.69 | 0.13 | p<0.01 |
| Avmax<br>(m/s)** | 0.79 | 0.16 | 0.85 | 0.16 | 0.70 | 0.11 | p<0.01 |
| E/A | 1.12 | 0.31 | 1.18 | 0.31 | 1.02 | 0.29 | NS |
| MR (m/s) | 5.62 | 0.52 | 5.62 | 0.52 |  |  |  |
| TR (m/s) | 2.42 | 0.53 | 2.42 | 0.53 |  |  |  |

Clinical and radiographic data of CMVD dogs at inclusion showed the following relevant findings: holosystolic mitral murmur (degree: 5% II, 29% III, 52% IV, 14% V), cough (28.5%), bronchial pulmonary pattern (48%), interstitial pulmonary pattern (5%), increased VHS (62%). Treatment for CHF was started at inclusion in 3 CMVD dogs with the triple therapy: benazepril 0.5 mg/kg PO BID, furosemide 2 ± 0.3 mg/kg PO BID and pimobendan 0.25 mg/kg PO BID. Healthy dogs showed normal VHS, tracheal position and caudal margin of the heart; while pulmonary pattern differ from trivial to sever diffuse bronchial pattern.

Follow up at 6-12-18 months is shown in Fig 1. Two CMVD dogs were lost during the study period and were censored for statistical analysis. At 6 month 11 CMVD dogs had WCD (2 of them with CHF) and no dogs had WRF. At 12 month, 14 CMVD dogs had WCD (5 of them experienced at least 1 episode of CHF), 3 of them with WRF. Four CMVD dogs showed stable kidney and heart function during the study period. At the end of the study period, 8 CMVD dogs experienced at least one episode of CHF and received triple therapy alone (n=4), triple therapy and spironolactone 2mg/kg PO SID (n=3) or triple therapy and anti-arrhythmic drugs (digoxin and diltiazem) (n=1). Two dogs of the healthy group at inclusion developed trivial mitral insufficiency and 3 developed proteinuria, 1 of them with low urine specific gravity = 1009.

**Fig 1.**
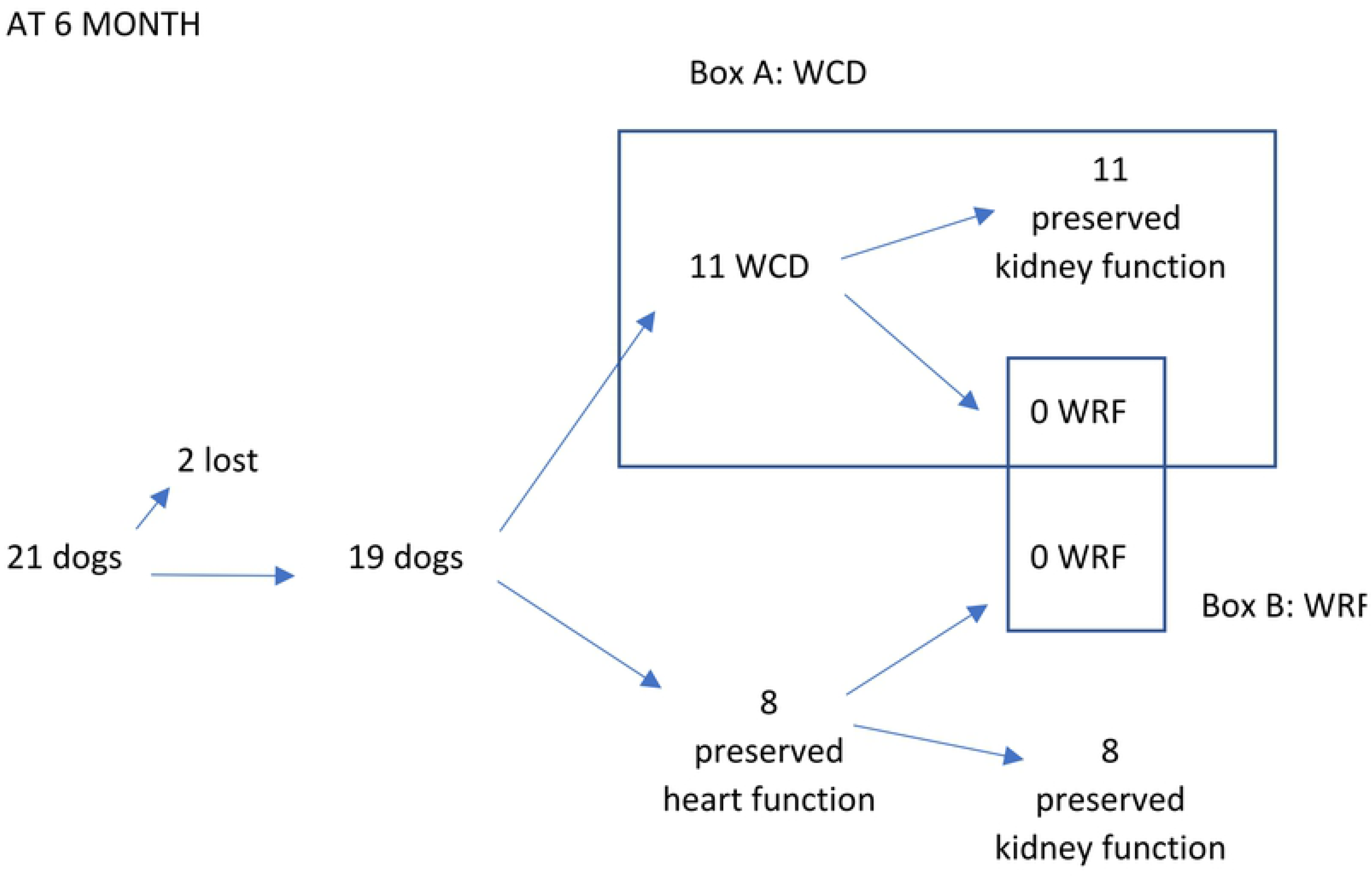
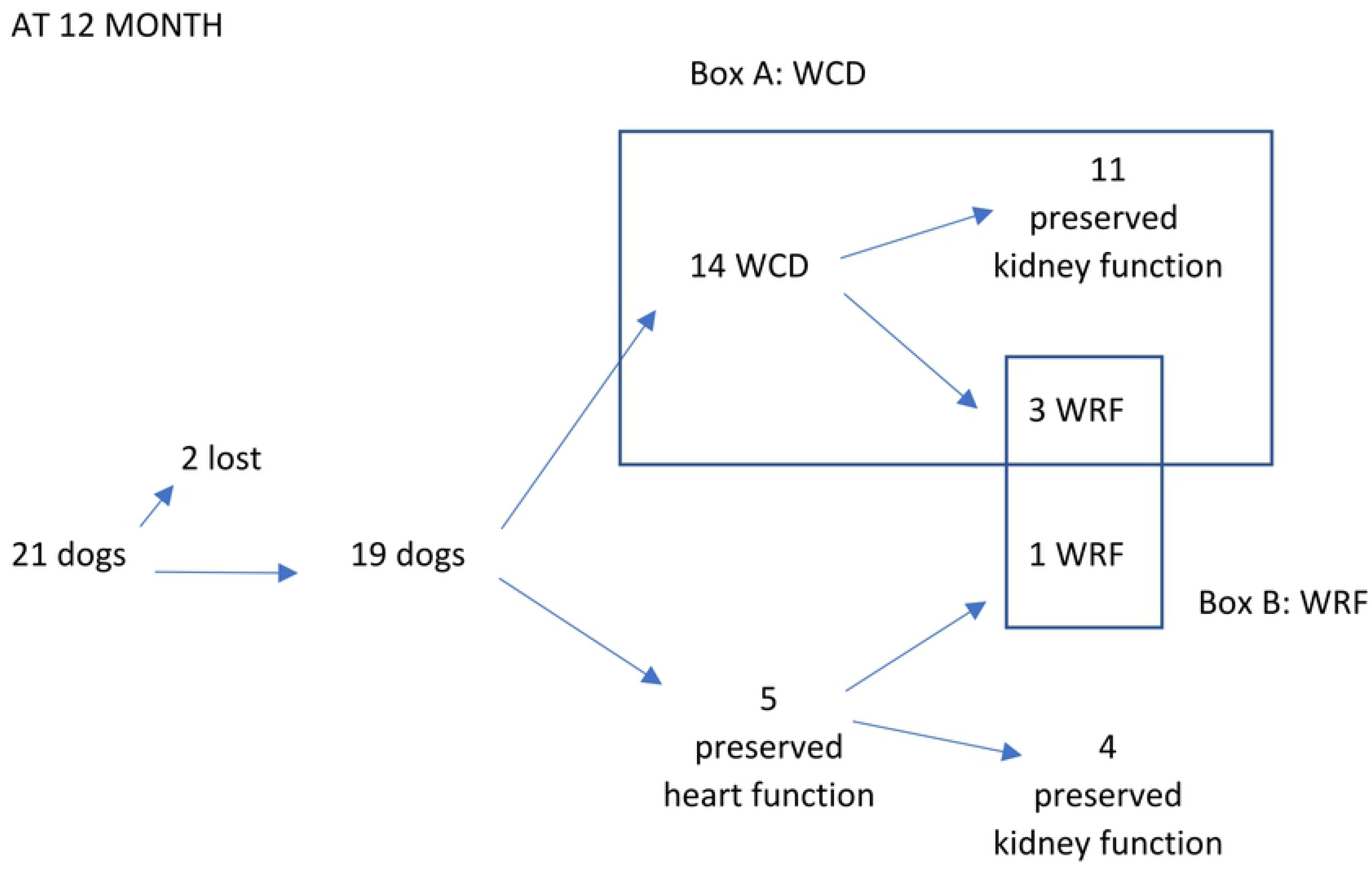
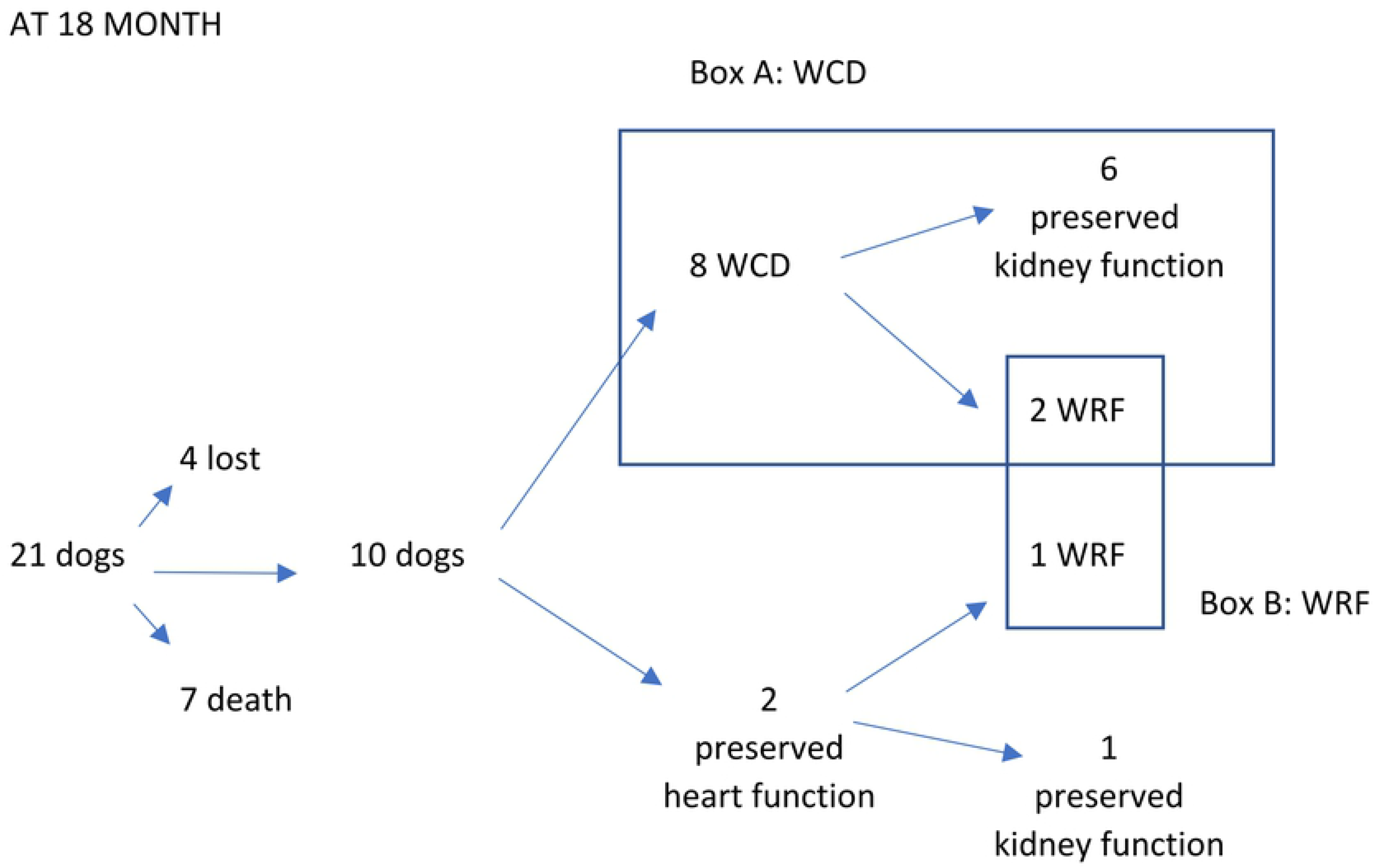
Follow up. Follow up at respectively 6 (A), 12 (B) and 18 (C) months of CMVD dogs and WCD/WRF status.

Considering the splitting of CMVD dogs into two groups according to Table 1 (dogs with/without WRF) there was not difference between the groups for the variables of WCD. Considering the splitting of dogs into two groups (dogs with/without WCD) there was no difference between the two groups for any variable of WRF.

At the end of the follow-up period (October 2014), 7 CMVD dogs died or were euthanized because of cardiac-related causes (n=2), renal-related causes (n=2) or other causes (n=3); 4 dogs of the healthy group at inclusion died or were euthanized because of non-cardiac and non-renal causes. Twelve CMVD dogs and 14 dogs of the healthy group were still alive, and 4 dogs (2 for each group) were lost at follow-up. Median survival time could not be calculated because <50% of the population died in the study period. In Fig 2 the KM curves are shown; however, due to the small sample size and to the fact that all healthy cases are censored, it is worthless to hazard statistical comparisons between survival times.

**Fig 2.**
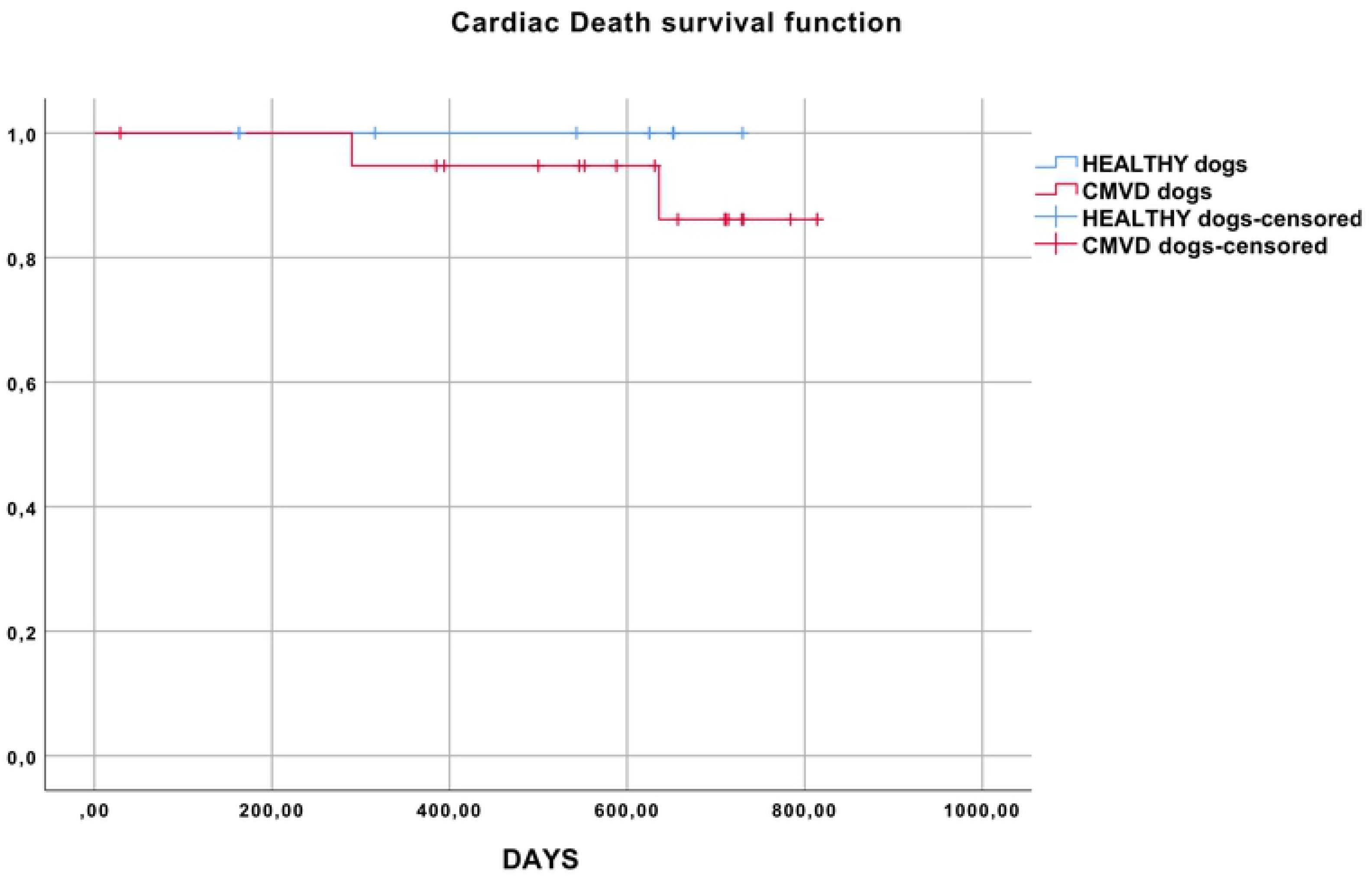
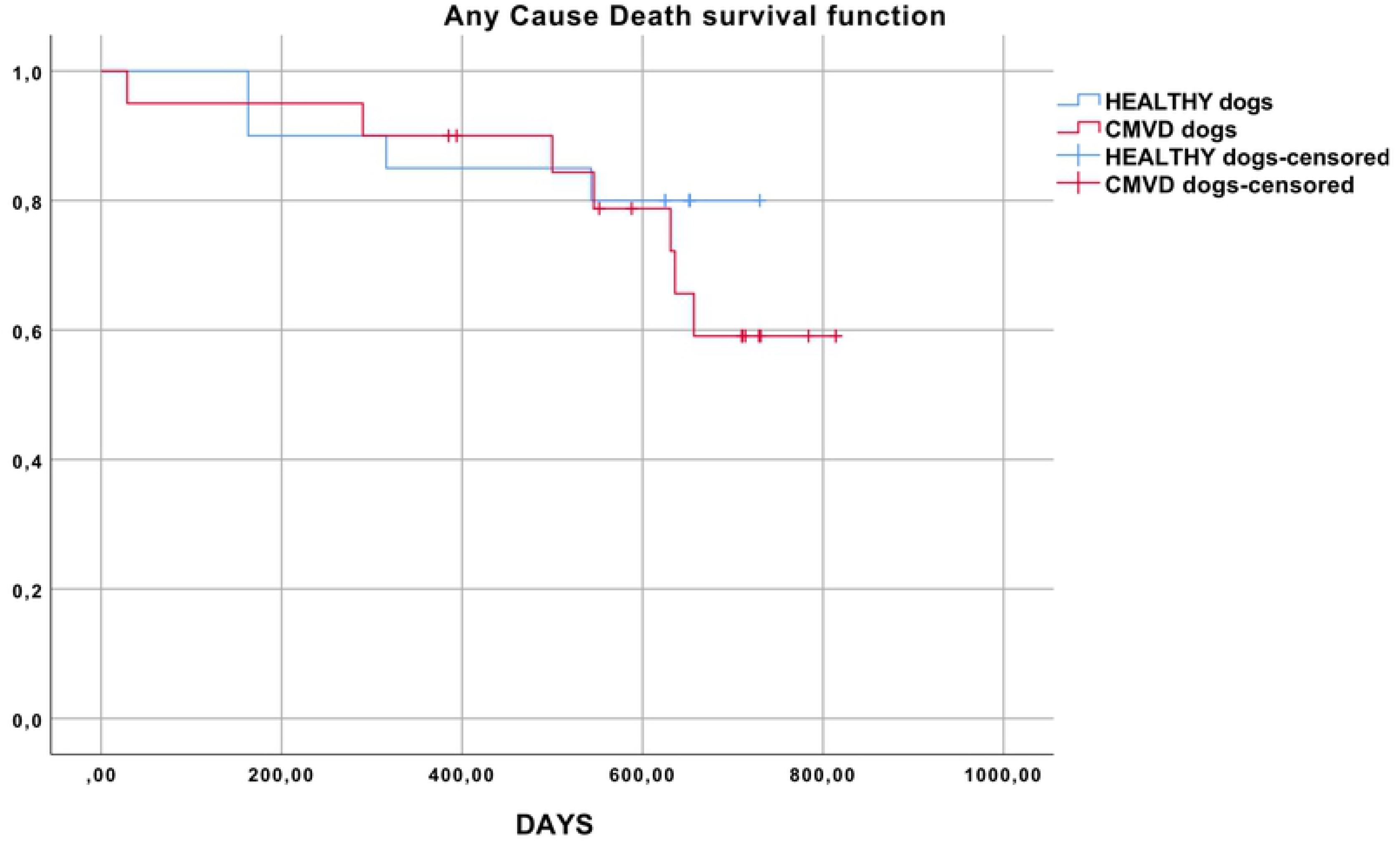
Survival curves. Kaplan-Meier survival curves for dogs with CMVD (red line) and healthy dogs (blue line) considering Cardiac Death (A) and Any cause Death (B).

Characteristics of the healthy and CMVD group and laboratory variables. White blood cell count (WBC), red blood cell count (RBC), hemoglobin (Hb), hematocrit (Ht), mean corpuscular volume (MCV), mean corpuscular hemoglobin (MCH), mean corpuscular hemoglobin concentration (MCHC), serum urea (UREA), serum creatinine (sCr), glucose (GLY), total proteins (TP), urinary proteins to urinary creatinine ratio (UPC), urine specific gravity (USG). Asterisks indicate statistically significant difference between healthy group and CMVD group:*p<0.05, **p<0.01.

Echocardiographic parameters in healthy and CMVD dogs and statistically significant difference between the two groups Heart rate (HR), left ventricular internal diameter (LVID) in diastole (d) and systole (s), normalized left ventricular end-diastolic diameter calculated according to Cornell’s method of allometric scaling (LVEDDn), Enddiastolic volume index (EDVI) Endsystolic volume index (ESVI), left ventricular ejection fraction (EF%), Fractional shortening (FS%), left atrial diameter (LA), Aortic root diameter (Ao), left atrial to aortic root ratio (LA/Ao), Mitral valve inflow (E peak velocity - EVmax, A peak velocity, E/A ratio), peak velocity of mitral and tricuspid regurgitations (MR and TR). Asterisks indicate statistically significant difference between healthy group and CMVD group:* p<0.05, **p<0.01.

## Discussion

Previous studies in veterinary medicine support the cardio-renal connection described in human medicine [8–12]. Our study assessed over time the most commonly used parameters of renal function (sCr, UREA, UPC and urine specific gravity), in 21 dogs affected by CMVD and 20 healthy dogs. The prevalence of azotemia in dogs affected by CMVD in this study (16%) is higher than the prevalence reported in the general population of dogs (0.05%-5.8%), but is lower than the prevalence reported in our retrospective study (25%) on a group of 158 dogs with CMVD [12, 30, 31]. Most importantly, we previously evidenced that an advanced ACVIM class could be predictive for advanced IRIS stage and *vice versa*, however the results of this study do not support a cardiorenal connection due to the lack of difference between the analyzed groups in developing WCD or WRF [12]. Based on the absolute number of cases experiencing WCD or WRF, we can assert that more dogs with WCD experience WRF than dogs without WCD, however, we are not able to clarify if these findings are secondary to the aged-related coexistence of CMVD and renal damage, more than cardio-renal syndrome. We didn’t find any correlation between diuretic therapy and renal function. We evidenced that one dog receiving triple therapy plus digoxin and diltiazem experienced multiple CHF episodes and developed azotemia, however, the essential link between severe CMVD and advanced therapy for medical management of heart disease (more severe CMVD require more aggressive therapy) made other consideration regarding therapy and renal function unreliable. Survival analysis was influenced by the absence of cardiac or renal death in the healthy group and didn’t add any useful information about the role of heart or kidney function on survival in dogs with CMVD. In fact, the 11% of healthy dogs at inclusion developed CMVD and none of them experienced cardiac related death. It has to been highlighted that low prevalence of CMVD in the control group is likely related to the difference of body weight and age. The CMVD group was representative of the general population of dog affected by CMVD (small and medium size breeds of middle to old age), while the healthy group included younger and heavier dogs that fulfilled the inclusion criteria. Because of the difference in life expectancy between small and large breed dogs, according to the human/pet analogy chart modified from Fortney WD, large breed dogs had to be consider “geriatric” when CMVD dogs were considered as “senior”. Hematocrit has been reported to be significantly lower in geriatric compared to senior dogs by Willems and colleagues; this finding could explain the difference in Ht between our two groups [32]. The 5% of control dogs at inclusion developed CKD stage IRIS 1, similarly to the prevalence of azotemia reported in the general population of dogs (5,8%), and none of them experienced renal related death [30].

The main limitations of this study are related to the small number of dogs included. However, the strict inclusion criteria allowed us to rule out some major confounding factors. The controls group did not match for age and weight with the study group and was not representative of the general population of dogs with CMVD, however this is a major problem of most of the studies about CMVD, due to its elevated incidence in the geriatric small breed dog population. Moreover, it has been reported that reference intervals of sCr should be based on BW categories; we established a cutoff of 1.4 mg/dl that could delay early diagnosis of renal dysfunction in small breed dogs and inversely might be too low for large and giant dogs [33]. The 11% of healthy dogs developed proteinuria; other causes of proteinuria (e.g. leishmaniosis, heartworms) were not evaluated in these dogs because of owner’s economics restraint. Symmetric dimethylarginine (SDMA) was not evaluated due to its late inclusion in IRIS guideline (2015).

## Conclusions

Congestive heart failure didn’t directly induced WRF in the studied population. Diuretics therapy, increasing in radiographic parameter VHS and/or increasing of the echocardiographic parameters elected as indicative of WCD, didn’t induce increasing sCr or appearance of azotemia and/or proteinuria. However, considering the prevalence of azotemia in the CMVD group (higher than the prevalence in the general population of dogs analyzed in previous paper), data suggests a link between heart and kidney function. We couldn’t exclude aged-related coexistence of CMVD and renal damage. A bigger number of dogs at inclusion is required to reach statistical significance.

## Acknowledgements

None

